# Genomic Diversity in *Porphyromonas*: Evidence of *P. catoniae* Commensality in Lungs

**DOI:** 10.1101/2024.11.09.622806

**Authors:** Lourdes Velo-Suarez, Yann Moalic, Charles-Antoine Guilloux, Jacky Ame, Claudie Lamoureux, Stéphanie Gouriou, Rozenn Le Berre, Clémence Beauruelle, Geneviève Héry-Arnaud

## Abstract

2.

Data on the genomics of *Porphyromonas* species other than *Porphyromonas gingivalis* (POTG), particularly within pulmonary environments, are scarce. In this study, we conducted whole genome sequencing (WGS) on pulmonary isolates of POTG, specifically *P. catoniae* (n=3), *P. pasteri* (n=1), and *P. uenonis* (n=2), from people with cystic fibrosis. These genomic analyses were complemented with antimicrobial susceptibility tests for these strains.

We compared the genomic sequences of these pulmonary isolates with those of previously characterized *Porphyromonas* species. Our study revealed a distinct clade differentiation between non-pigmented and pigmented *Porphyromonas* species. Interestingly, unlike *P. gingivalis*, the pulmonary POTG strains lacked known virulence genes, except a putative hemolysin gene. Regarding antibiotic resistance, notable resistances were limited to vancomycin in *P. catoniae* and clindamycin in *P. uenonis*.

These findings support the hypothesis that POTG species may predominantly behave as commensals in the lung environment rather than as pathogens.

**Impact statement:** *Porphyromonas* constitutes a prevalent genus of anaerobic bacteria within the respiratory tract. Despite its ubiquity, the precise taxonomic delineation of *Porphyromonas* species implicated in respiratory conditions needs to be better defined, as most microbiome analyses report results only at the genus level. Consequently, data about these pulmonary species often default to associations with *Porphyromonas gingivalis*, a well-characterized oral pathogen. In this study, we sequenced the complete genomes of six *Porphyromonas* strains isolated from the airway microbiota of people with cystic fibrosis (CF) to enhance the representation and identification of *Porphyromonas* other than *gingivalis* (POTG) species. Our phylogenomic analysis aimed to elucidate the diversity within pulmonary POTG. Notably, genomes of the commensal species *Porphyromonas catoniae* isolated from CF patients did not harbor virulence genes typically associated with *P. gingivalis*. Additionally, phenotypic resistance profiling against nine clinically relevant antibiotics revealed low resistance levels, notwithstanding the frequent antibiotic treatments administered to CF patients. Our findings provide compelling evidence for the non-pathogenic role of these POTG species in the pulmonary environment.

**Data summary:** Six *Porphyromonas* genomes generated in this study are available in the Sequence Read Archive and GenBank databases under BioProject accession PRJEB75658.

The raw data set generated during the current study is available in the European Nucleotide Archive (repository with the project accession number.

The authors confirm that all supporting data, code, and protocols have been provided within the article or through supplementary data files.

## 5. Introduction

In recent years, the composition of the pulmonary microbiota has garnered significant interest, particularly regarding its role in chronic respiratory diseases. The complex ecology of the lung microbiota, rich even in anaerobes among healthy populations, provides a pivotal lens through which disease progression can be viewed (1-15). *Porphyromonas*, ranking among the top ten anaerobic taxa in healthy lungs, exemplifies the metabolic diversity of lung anaerobes, spanning both commensal and pathogenic roles (16, 17). Notably, *Porphyromonas gingivalis*, a key pathogenic species, has been extensively studied for its virulence and adaptation strategies. However, other *Porphyromonas* species, such as *Porphyromonas catoniae* and *Porphyromonas pasteri*, typically associated with the healthy oral microbiota, have yet to be investigated, particularly from pulmonary sources (18, 19).

Our study aims to fill this gap by performing whole-genome sequencing (WGS) on pulmonary *Porphyromonas* isolates previously collected (17). We explore the genomic distinctions between oral and pulmonary strains and identify potential genes of interest within the genomes of pulmonary *Porphyromonas* other than *gingivalis* (POTG). Additionally, we examine the antibiotic resistance genes (ARGs) in conjunction with phenotypic resistance profiles to assess whether POTG could act as a commensal species within the respiratory tract. Our findings could significantly impact our understanding of the potential commensal or pathogenic roles of less-studied *Porphyromonas* species.

## 6. Methods

### Isolation and Culture of Strains

The clinical strains of POTG employed in this study comprised two strains of *P. catoniae* (PC), one strain of *P. pasteri* (PP), and two strains of *P. uenonis* (PU), all isolated from the sputum of people with cystic fibrosis (CF). Sample selection adhered to the Murray-Washington quality score to exclude contamination from saliva (20). Collection was conducted using a device specifically designed to preserve an anaerobic environment as previously described (17). This involved plating the samples within an anaerobic chamber (Bactron 300, Sheldon Manufacturing) that maintained a gas composition of 5% CO_2_, 5% H_2_, and 90% N_2_.

The bacterial colonies were initially cultivated on Anaerobe Basal Agar supplemented with 5% sheep blood (ABA-SB) from ThermoFisher. ABA-SB media containing kanamycin-vancomycin and colistin-nalidixic acid were also employed for selective cultivation. The strains were identified using matrix-assisted laser desorption/ionization time-of-flight mass spectrometry (MALDI-TOF MS) with the Biotyper MBT system (Bruker). Additional confirmation for some colonies was achieved by 16S rRNA sequencing performed on an Applied Biosystems 3130xl instrument (ThermoFisher) using the BigDye Terminator Cycle Sequencing Kit Ready Reaction v3.1 (Applied Biosystems) as outlined by Lamoureux et al. (17).

*Porphyromonas* cultures were grown on ABA-SB plates for 4 to 7 days, followed by subculturing in a homemade Peptone Yeast (PY) liquid medium for genomic DNA extraction. The cultures were incubated for an additional three days at 37°C within the anaerobic chamber, ensuring optimal growth conditions for subsequent analyses.

### DNA Extraction and Sequencing

DNA was extracted using the QIAamp DNA Mini Kit (Qiagen), following the manufacturer’s instructions, with elution in 100 μL. We employed two distinct sequencing platforms to ensure comprehensive genomic data coverage:

#### PacBio Sequel System

Library preparation was carried out using the Template Prep Kit V2 per the manufacturer’s specifications. Sequencing was conducted in a single run at the Faculty of Medicine, Brest, France, using the Sequel Sequencing Kit 3.0 and the Sequel SMRT Cell 1M v3 LR Tray. This system is noted for its long-read capabilities, crucial for assembling complex genomic regions.

#### Illumina MiSeq System

Libraries were prepared using the TruSeq Nano kit, following the manufacturer’s guidelines. Whole genome sequencing was performed in a single run using V3 chemistry (2×250 bp) at the Genotoul platform, Castanet-Tolosan-Auzeville, France. This system is favored for its high-precision short-read sequencing.

This hybrid sequencing strategy was designed to maximize the coverage and accuracy of the genomic data, facilitating a deeper understanding of the genetic intricacies of the *Porphyromonas* species under study.

### Genome Assembly

#### Quality Control

Raw sequence reads underwent rigorous quality control to assess and filter out low-quality sequences. Adapter sequences and substandard reads were removed using bbduk from the bbmap suite v38.92-0, setting a quality base threshold of 20 and a minimum length requirement of 200 bp. Further, we assessed quality and potential contamination using kat v2.4.2 and kraken2 v2.1.2 to identify human and other non-target sequences.

#### Genome Assembly and Annotation

Hybrid genome assemblies were conducted using the Unicycler pipeline v0.4.8-beta, incorporating a suite of tools including spades.py v3.13.0, racon v1.3.3, makeblastdb v2.9.0+, tblastn v2.9.0+, bowtie2-build v2.3.5, bowtie2 v2.3.5, samtools v1.9, java v11.0.1, and pilon v1.23 (21). Assembly quality statistics were generated using Quast v5.02, and genome completeness and contamination levels were evaluated with CheckM v1.0.18.

#### Scaffold Profiling and Gene Identification

We utilized Anvi’o v7 for scaffold profiling, employing Prodigal v2.6.354 with default settings for gene identification and HMMER v3.355 to detect bacterial single-copy genes and ribosomal RNA-based Hidden Markov Model (HMM). Short reads from each strain were mapped back to their respective scaffolds using Bowtie2 v2.4.4-0, with a minimum identity threshold of 95%. The mapped reads were then stored as BAM files using samtools. Anvi’o was subsequently used to profile each BAM file, allowing for precise estimation of coverage and detection statistics for each scaffold.

### Phylogenomics and Pan-genome Analysis

To determine the phylogenetic position of our six pulmonary *Porphyromonas* strains, we accessed 99 assigned *Porphyromonas* genomes and 23 non-redundant *Alloprevotella* genomes (serving as outgroup) from the NCBI database using the ncbi-genome-download tool (https://github.com/kblin/ncbi-genome-download). We assessed whole-genome similarity using the Average Nucleotide Identity (ANI), computed with FastANI v1.32. For phylogenomic and pangenomic analyses, only genome pairs from NCBI exhibiting FastANI values greater than 98% were retained, narrowing the dataset to 59 *Porphyromonas* genomes and 15 outgroup genomes. Using HMMER v3.3, we extracted HMM hits for each genome, focusing on 36 ribosomal single-copy core genes. These genes were aligned, and gaps affecting more than 50% of nucleotide positions were removed using trimAl v1.4.1, ensuring alignment quality for subsequent analyses. The phylogenetic analysis of these high-quality ribosomal gene alignments was conducted on a concatenated dataset. Substitution models for each gene were automatically selected from the GTR + F + Γ + I + R family using model-testing algorithms to find the best-fitting partition scheme. Phylogenetic trees were inferred using maximum likelihood estimation with proportional variation in branch lengths across partition subsets, implemented in IQ-TREE2 v1.7, using the LG + F + R4 model. The robustness of the phylogenetic tree was supported by 5,000 bootstrap replicates, providing statistical confidence in the branching patterns observed.

To explore the genetic diversity and evolutionary relationships within the *Porphyromonas* genus, we employed the Anvi’o platform to compute and visualize the pan-genome of *Porphyromonas*, encompassing both our strains and reference genomes. We stored 65 genomes (59 reference genomes from NCBI and our 6 pulmonary strains) in an Anvi’o database using the ‘anvi-gen-genomes-storage’ command. For the pan-genomic analysis, we focused on closely related genomes available from NCBI. Creating the pan-genome involved passing the stored database to the ‘anvi-pan-genome’ command, which utilized NCBI’s BLAST to assess gene similarity within and between the genomes. To cluster these genes into functionally coherent groups, we applied the Markov Cluster Algorithm (MCL), facilitating the identification of core and accessory genomic components across the studied *Porphyromonas* species. This comprehensive analysis allows us to better understand the genomic architecture and variability among species, which is crucial for elucidating their evolutionary trajectories and potential functional roles in different environmental niches.

### Genome Annotation and Characterization of Pulmonary *P. catoniae* Strains

The assembled genomes of *P. catoniae* isolated from pulmonary sources were comprehensively annotated and analyzed using several advanced bioinformatics tools. We utilized Prokka v1.13 (22) for rapid annotation, the MicroScope Microbial Genome Annotation and Analysis Platform (MaGe) (23), the NCBI integrated Prokaryotic Genome Annotation Pipeline (PGAP), and the RAST server v2.0 (Overbeck, 2014). These tools were employed with default parameters to ensure a standardized approach across all annotations.

To delve deeper into the metabolic capabilities of our lung *P. catoniae* strains, we compared their profiles with those of *P. catoniae* ATCC 51270 and several strains of *P. gingivalis* (ATCC 33277, TDC60 and W83), using the KEGG and BioCyc databases. This comparison was facilitated by the analytical tools available on the MicroScope platform, which also enabled the detection of macromolecular systems via MacSyFinder v1.0.2, prediction of CRISPR arrays using CRISPRCasFinder v4.2.19, identification of antibiotic resistance genes through the CARD database v3.0.2, and exploration of secondary metabolite regions using antiSMASH v5.0.0. Additionally, we investigated the presence of mobile genetic elements in the genomes using Phaster, viralVerify (24), and VirSorter2 (25). These analyses provided insights into the genomic flexibility and potential adaptative mechanisms of *P. catoniae* in the pulmonary environment.

### Phenotypic Tolerance and Antibiotic Resistance Testing

Given the genomic identity established among the strains PC103, PC123, and PC135, phenotypic tests were exclusively conducted on the PC103 strain to evaluate its environmental tolerance capabilities.

#### Oxygen Tolerance

The PC103 strain’s oxygen tolerance was assessed by suspending the bacteria in non-oxygen-reduced physiological serum at two initial concentrations: a high concentration of 10^7^ CFU.mL^-1^ and a lower concentration of 10^3^ CFU.mL^-1^. The suspensions were exposed to ambient oxygen, and samples were periodically collected, plated, and incubated for 7 days at 37°C in an anaerobic chamber (5% CO_2_, 5% H_2_, 90% N_2_) to measure the residual inoculum.

#### Acid Tolerance

PC103’s acid tolerance was tested using ABA-SB medium acidified with 1M HCl to achieve pH values of 6.86, 6.38, 6.0, 5.5, and 5.0. Actively growing cultures of *P. catoniae* PC103 on standard ABA-SB were transferred to the pH-adjusted ABA-SB. These plates were incubated under the same anaerobic conditions for 7 days, after which colony growth was evaluated.

#### Cadmium Tolerance

Cadmium tolerance was examined using ABA-SB medium supplemented with increasing cadmium concentrations, ranging from 0 to 80 mg.L^-1^. Cultures of PC103 grown on standard ABA-SB were subcultured onto cadmium-enriched ABA-SB and incubated for 7 days in an anaerobic environment at 37°C. Growth assessment was performed by examining the colonies’ presence. These tests provide valuable insights into the robustness of the PC103 strain under various stress conditions, which may contribute to understanding its survivability and adaptability in the pulmonary niche.

#### Antibiotic resistance

The antibiotic resistance profiles of six *Porphyromonas* strains isolated from pulmonary sources were assessed following the 2020 guidelines provided by the European Committee on Antimicrobial Susceptibility Testing (EUCAST) and the Committee for Antibiogram of the French Society of Microbiology (CA-SFM) (26). Freshly isolated cultures from ABA-SB were suspended to a density equivalent to 1 McFarland standard in Anaerobe Basal Broth (ABB) (ThermoFisher). These suspensions were then plated onto ABA-SB agar to establish a uniform bacterial lawn. The antibiotics tested included amoxicillin (25 µg), amoxicillin/clavulanic acid (20 µg/10 µg), clindamycin (2 µg), imipenem (10 µg), metronidazole (5 µg), moxifloxacin (5 µg), piperacillin (30 µg), and piperacillin/tazobactam (30 µg/6 µg) (ThermoFisher). Additionally, a vancomycin disk (30 µg) was included due to the noted susceptibility of other *Porphyromonas* species, although *P. catoniae* typically shows resistance (27). The inoculated agar plates were incubated in an anaerobic chamber maintained at 37°C for at least 48 hours to allow adequate growth and interaction between the bacterial isolates and the antibiotic agents.

## 7. Results

### Whole genome sequencing (WGS) of POTG (*P. catoniae, P. pasteri*, and *P. uenonis*)

The genomic characteristics of pulmonary strains of the *Porphyromonas* genus, specifically *P. catoniae, P. pasteri*, and *P. uenonis*, were investigated through WGS. The GC% content of these strains fell within the range of 51% to 56%, consistent with prior findings for this genus. Utilizing *de novo* hybrid assembly methods, the genomes of the pulmonary POTG were effectively reconstructed, resulting in contig numbers ranging from 1 to 66, with N50 values spanning from 216,372 bp to 2,401,504 bp. The largest contig length observed for *Porphyromonas* ranged from 320,622 bp to 2,401,504 bp. Notably, the *de novo* assembly lengths agreed with previously reported median total lengths for each species. Assessment of genome completeness revealed values ranging from 95.05% to 99.92%. Detailed genomic statistics are provided in Table 1.

**Table 1.** Sequencing and assembly statistics of pulmonary strains of *Porphyromonas* other than *P. gingivalis* (POTG)

| Strain | POTG species | Unicycler assembly | N50 | Largest contig | GC% | Contigs number | Completeness | Contamination | Predicted CDS |
| --- | --- | --- | --- | --- | --- | --- | --- | --- | --- |
| PC103 | *P. catoniae* | 2,032,211 | 2,032,211 | 2,032,211 | 51.00 | 1 | 95.05 | 0.157 | 1619 |
| PC123 | *P. catoniae* | 2,032,214 | 1,601,551 | 1,601,551 | 51.00 | 3 | 95.05 | 0.157 | 1622 |
| PC135 | *P. catoniae* | 2,032,218 | 2,032,218 | 2,032,218 | 51.09 | 1 | 95.05 | 0.157 | 1620 |
| PP88 | *P. pasteri* | 2,225,553 | 2,205,487 | 2,205,487 | 55.75 | 2 | 98.87 | 0 | 1703 |
| PU103 | *P. uenonis* | 2,401,504 | 2,401,504 | 2,401,504 | 52.31 | 1 | 99.92 | 0 | 1882 |
| PU61 | *P. uenonis* | 2,325,647 | 216,372 | 320,622 | 52.49 | 66 | 99.92 | 0 | 1876 |

### Analysis of Average Nucleotide Identity (ANI) in Pulmonary *Porphyromonas* Strains

ANI analysis of pulmonary strains revealed two distinct genetic clusters among the *Porphyromonas* species (Table 2). Non-pigmented strains of *P. catoniae* and *P. pasteri* formed one group, while all *P. uenonis* strains comprised the second group. Specifically, strains PC103, PC123, and PC135 were identified as belonging to the same species, adopting a 95% similarity threshold for species delineation, as established by Konstantinidis and Tiedje (28) and Konstantinidis *et al*. (29). These strains exhibited ANI values of 96.6% and 96.7% with the reference strain of *P. catoniae* (F0037).

**Table 2.** Average nucleotide identity between *Porphyromonas* genomes Phylogenomic Analysis.

| Strain* | PC103 | PC123 | PC | PC135 | PP88 | PP | PU103 | PU61 | PU | PA | PB | PE | PS | PG |
| --- | --- | --- | --- | --- | --- | --- | --- | --- | --- | --- | --- | --- | --- | --- |
| PC103 | 100 | 99.9 | 96.6 | 99.9 | 79 | 78.5 | <80 | <80 | <80 | <80 | <80 | <80 | <80 | <80 |
| PC123 | 99.9 | 100 | 96.7 | 99.9 | 78.8 | 78.4 | <80 | <80 | <80 | <80 | <80 | <80 | <80 | <80 |
| PC135 | 99.9 | 99.9 | 96.6 | 100 | 79 | 78.6 | <80 | <80 | <80 | <80 | <80 | <80 | <80 | <80 |
| PP88 | 79 | 78.8 | 78.7 | 79 | 100 | 96.4 | <80 | <80 | <80 | <80 | <80 | <80 | <80 | <80 |
| PU103 | <80 | <80 | <80 | <80 | <80 | <80 | 100 | 97.6 | 97.4 | 88.8 | <80 | <80 | <80 | <80 |
| PU61 | <80 | <80 | <80 | <80 | <80 | <80 | 97.6 | 100 | 97.4 | 88.9 | <80 | <80 | <80 | <80 |

\*PC, *Porphyromonas catoniae* F0037; PP, *Porphyromonas pasteri* JCM 30531; PU, *Porphyromonas uenonis* DSM 23387; PA, *Porphyromonas asaccharolytica* DSM 20707; PB, *Porphyromonas bennonis* DSM 23058; PE, *Porphyromonas endodontalis* 52278C01; PS, *Porphyromonas somerae* DSM 23386; PG, *Porphyromonas gingivalis* ATCC 33277

Conversely, strain PP88 showed only 78.7% to 79% similarity with strains PC103, PC123, and PC135, indicating significant genomic divergence. However, PP88 displayed a higher similarity of 96.4% with the reference strain of *P. pasteri* (JCM 30531), suggesting a closer genetic relationship with this species. Furthermore, strains PU61 and PU103 showed a high similarity of 97.6%. These strains and the reference strain of *P. uenonis* (DSM 23387), which was initially isolated from a human sacral decubitus ulcer, shared a similarity of 97.4% ANI. Notably, all analyzed POTG lung strains demonstrated less than 80% similarity with the reference strain of *P. gingivalis* (ATCC 33277), further highlighting the genetic distinctiveness of the pulmonary strains within this group.

### Phylogenomic Analysis

In the phylogenomic analysis, ribosomal sequences from all publicly available *Porphyromonas* strains were concatenated to construct a comprehensive phylogenomic tree (Figure 1). This analysis enabled the clear differentiation of species within the *Porphyromonas* genus. The non-pigmented strains of the POTG, including *P. pasteri* and *P. catoniae*, formed a distinct clade, demonstrating their phylogenetic separation from other pigmented *Porphyromonas* species.

**Fig 1.**
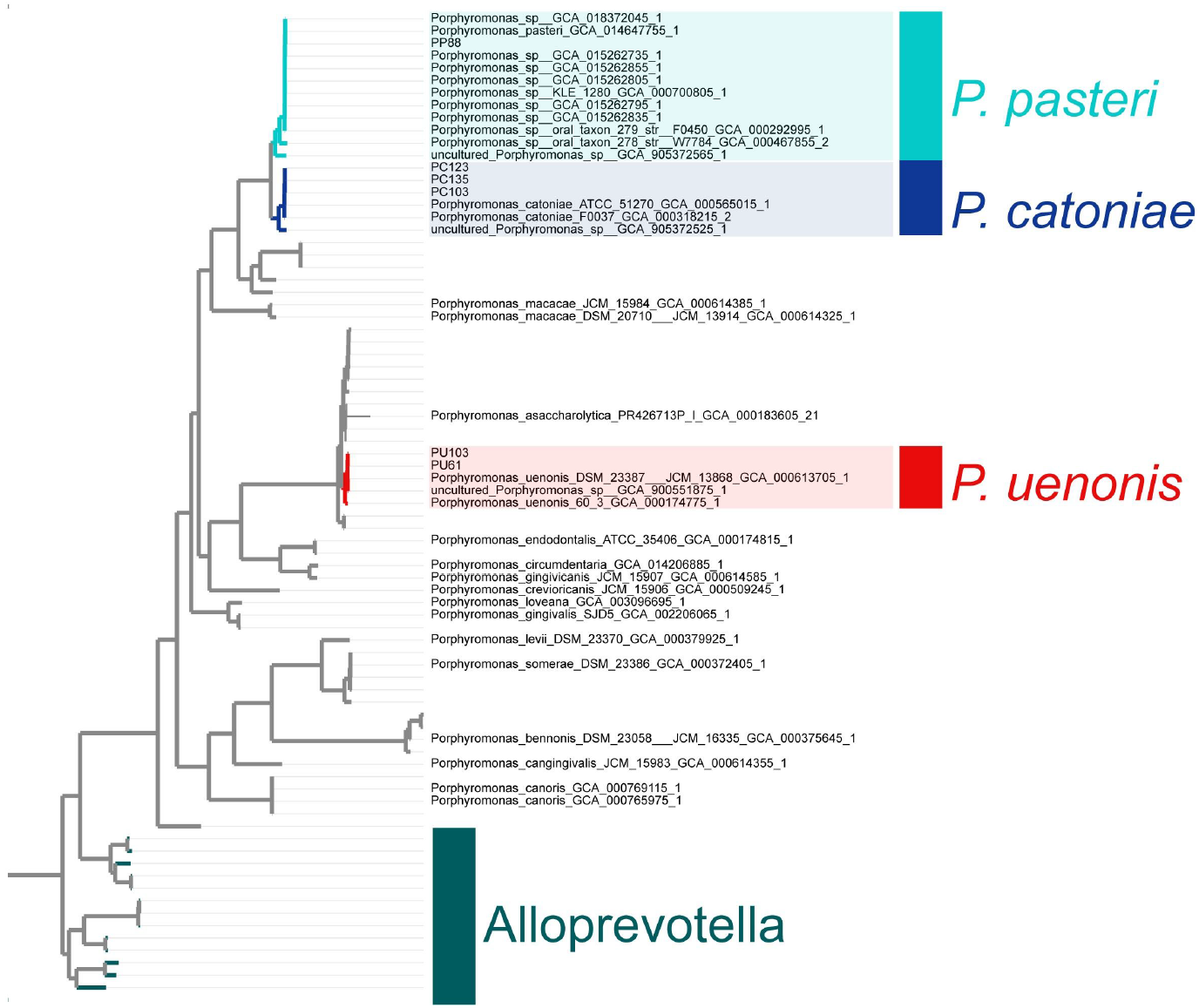
Phylogenomic tree of *Porphyromonas* pulmonary species. Tree based on concatenating ribosomal sequences using IQ-TREE natively with ModelFinder Plus (30).

In detail, lung strains PU61 and PU103 closely aligned with all reference genomes of *P. uenonis*, indicating strong phylogenetic consistency within this species. Similarly, PC103, PC123, and PC135 clustered closely with all reference genomes of *P. catoniae*, confirming their classification within this species. Additionally, PP88 was phylogenetically aligned with all reference genomes of *P. pasteri*, underscoring its species-specific grouping.

### Pan-Genomic Analysis of *Porphyromonas* Strains

A comprehensive pan-genomic analysis was conducted on lung strains of *Porphyromonas* from our collection alongside reference genomes from the POTG group. This analysis revealed a core genome consistent across all *Porphyromonas* species, comprising 357 gene clusters encompassing 8,587 open reading frames (ORFs). A species-specific accessory genome was identified exclusively in *P. catoniae*, containing 255 gene clusters (419 ORFs). The study aimed to elucidate potential virulence factors among the POTG, mainly focusing on non-pigmented *Porphyromonas* strains. The virulence genes examined were primarily those previously identified in *P. gingivalis* ATCC 33277 by Mendez *et al*. (31), who extensively searched for such genes in that strain. However, our current analysis did not detect genes associated with gingipains or hemagglutinin in the non-pigmented *Porphyromonas* genomes. Only one putative hemolysin gene was found, and an additional gene was identified as hyaluronidase/collagenase (K01197; EC: 3.2.1.35), indicating a reduced virulence gene profile in these strains. Regarding secretion systems, a type IX secretion system was detected in both PC103 and *P. catoniae* ATCC 51270 genomes, albeit incomplete, and a type IV secretion system was noted in *P. catoniae* ATCC 51270, also incomplete. Interestingly, specific metabolic pathways and transport mechanisms, such as the pentose phosphate pathway and iron transport genes, were explicitly identified in the lung strains of *P. catoniae*, suggesting adaptations to their niche. These screened genes and their potential implications are summarized in Table in Supplementary Information.

### Genome comparison of *Porphyromonas* strains

The comprehensive genomic analysis of the lung-derived *P. catoniae* strain PC103 reveals a single circular chromosome measuring 2,032,211 base pairs with a genomic DNA G+C content of 51.09%. Genome annotation, performed using the MaGe platform, predicted 1,669 genes, including 1609 protein-coding sequences (CDSs), as illustrated in Figure 3. The analysis also identified one CRISPR locus between 805,723 and 806,843 bp downstream of a CAS-type II cluster spanning from 800,799 to 805,615 bp. Additionally, two incomplete prophage regions were detected, each approximately 27 kb and 26 kb in size, showing 40% completeness.

The metabolic profile of *P. catoniae* PC103 was compared against *P. catoniae* ATCC 51270 and several *P. gingivalis* strains (ATCC 33277, TDC60, and W83) using the MicroScope platform. This comparison highlighted significant metabolic distinctions between the *P. catoniae* and *P. gingivalis* strains, irrespective of their source (oral or pulmonary). Key pathway differences are summarized in Table 3, including a cadmium transport system unique to *P. catoniae* strains and absent in *P. gingivalis* strains. Conversely, genes involved in protein citrullination were exclusive to *P. gingivalis* strains. *P. catoniae* strains exhibited unique sulfur metabolism pathways (sulfide oxidation and sulfur reduction) and an arginine-dependent acid resistance gene. The genomes also showed a partial presence of pathways for detoxifying oxidized GTP and dGTP and the degradation of superoxide radicals. Furthermore, the fatty acid biosynthesis pathways showed 60% completion for initiation and 80% completion for elongation processes. Additionally, various vitamin biosynthesis pathways were identified in *P. catoniae*, including those for folate, lipoic acid, pantothenate, riboflavin, and pyridoxal, contrasting with the degradation pathways for amino acids and sugars found in *P. gingivalis* genomes.

**Table 3.**
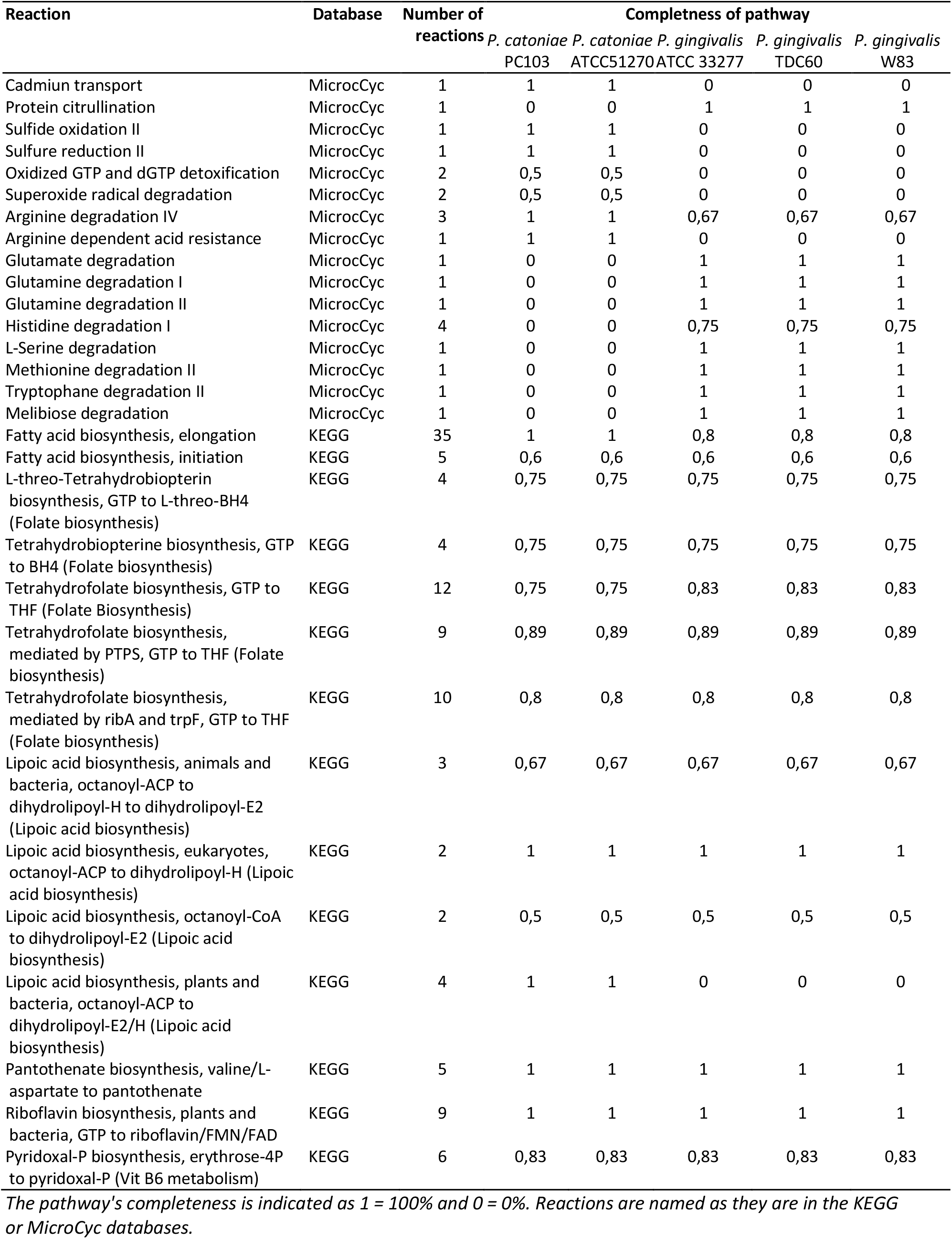
Metabolic comparison of *P. catoniae* PC103, *P. catoniae* ATCC 51270, *P. gingivalis* ATCC 33277, *P. gingivalis* TDC60, and *P. gingivalis* W83 with the MicroScope platform using KEGG and MicroCyc databases.

### *In Vitro* Confirmation of Genotypic Characteristics of *Porphyromonas* Strains

#### Oxygen Tolerance

The presence of genes related to the detoxification of oxidized GTP and dGTP and the degradation of superoxide radicals in *P. catoniae* PC103 suggested potential oxygen tolerance. To verify this, we exposed a suspension of PC103 to atmospheric oxygen concentrations, monitoring the bacterial survival over time. The results, depicted in Figures 3A and B, showed only a 1.86 log decrease in CFU per milliliter after 8 hours at an initial concentration of 10^7^ CFU.mL^-1^ and a 1.25 log decrease after 6 hours at 10^3^ CFU.mL^-1^.

#### pH Tolerance

The *in silico-identified* arginine-dependent acid resistance gene prompted an *in vitro* assessment of PC103’s acid tolerance. Cultures were grown on ABA-SB medium at various pH levels (6.86, 6.38, 6.0, 5.5, and 5.0) and incubated for 7 days. PC103 was able to grow at pH levels down to 5.5, but no growth was observed at pH 5.0, even with extended incubation.

#### Cadmium Resistance

The WGS of PC103 had revealed a cadmium transport gene, suggesting possible resistance to cadmium. This was tested by culturing PC103 on an ABA-SB medium supplemented with increasing cadmium concentrations (0 to 80 mg.L^-1^). The results demonstrated that PC103 could tolerate cadmium concentrations up to 20 mg.L^-1^, but growth was inhibited at 40 mg.L^-1^ and above.

#### Antibiotic Resistance

Investigations into the antibiotic susceptibility of pulmonary *Porphyromonas* species revealed sensitivity to a broad spectrum of antibiotics, with consistent antibiogram profiles across the species, except for specific resistances. Strains of *P. uenonis* (n=2) displayed resistance to clindamycin. Strains of *P. catoniae* (n=2), including PC103, were resistant to vancomycin. Intriguingly, no vancomycin resistance genes were identified in the reconstructed genomes of these strains using the CARD database, indicating a potential non-genomic resistance mechanism or an unidentified resistance gene.

## 8. Discussion

In this study, we conducted WGS of pulmonary strains of the POTG group, specifically *P. catoniae, P. pasteri*, and *P. uenonis*, isolated from the sputum of people with CF. Our genomic analysis revealed high nucleotide similarity among these strains and identified a distinct clade of non-pigmented *Porphyromonas* in the phylogenomic tree. Notably, *P. catoniae* was found to lack the virulence genes typical of *P. gingivalis*, except for a putative hemolysin gene.

*As reported in several studies, P. gingivalis* is a well-documented pathogen extensively associated with periodontitis (18, 32-34). Recent research has also implicated this bacterium in systemic diseases, including lung cancer (35), rheumatoid arthritis (RA) (36), and Alzheimer’s disease (37), highlighting its potential role beyond oral health. In contrast, the genus *Porphyromonas* has been detected in both diseased and healthy pulmonary niches using molecular and cultural methods (2, 5, 13-15, 17, 38). Specifically, targeted metagenomic analyses have frequently identified POTG strains, predominantly non-pigmented *Porphyromonas*, in pulmonary samples (2, 16, 17). Unlike *P. gingivalis*, species such as *P. catoniae* and *P. pasteri* are primarily associated with healthy individuals, suggesting potentially different roles in the microbial ecology of the lung (18, 39-41).

Historically, genomic data on non-pigmented *Porphyromonas* species have been sparse. A notable study by O’Flynn *et al*. (42) included a genome of *P. catoniae* isolated from a human gingival crevice, revealing significant genomic distinctions between *P. catoniae* and *P. gingivalis*. Key differences highlighted were the absence of hemagglutinin and gingipains, virulence factors in *P. gingivalis*, whereas genes for hemolysins and iron transport were detected in *P. catoniae*. Consistent with these findings, our current analysis of pulmonary non-pigmented *Porphyromonas* strains identified similar absences and noted the presence of a putative hemolysin (cds 1638831 1640078 - HANLCC_06880 hemolysin SO:0001217, UniRef: UniRef50_A0A0A2G0A3, UniRef: UniRef90_L1N9Y2).

Moreover, O’Flynn et al. reported the absence of glutamate enzymes in *P. catoniae* (42), except for a NADP-specific glutamate dehydrogenase (EC 1.4.1.4), which we also identified in our study, indicating its conservation across different isolates of this species. Interestingly, our research further established the presence of genes encoding pentose phosphate pathway enzymes in *P. catoniae*, contrasting with their absence in *P. gingivalis* genomes. This study also uncovered genes involved in synthesizing glutamate from ammonia and maintaining the glutamate pool in *P. catoniae*, elements missing in *P. gingivalis*. Conversely, genes related to the synthesis of ammonia from nitrate, the synthesis of vitamin B12 from uroporphyrinogen, and the release of iron from blood cells, typically found in *P. gingivalis*, were absent in *P. catoniae* (42).

Significantly, none of the *P. catoniae* genomes, including lung clinical strains and *P. catoniae* ATCC 51270, exhibited genes involved in protein citrullination, a process linked to the production of citrullinated protein antigens, which are specific markers for RA (43). This absence is crucial because *P. gingivalis* has been implicated in RA due to its ability to produce such antigens (44). The lack of citrullination genes in non-pigmented *Porphyromonas* suggests that these species play no role in RA and possibly other autoimmune diseases. This genetic profile supports the hypothesis that non-pigmented *Porphyromonas*, particularly *P. catoniae*, are less likely to contribute to autoimmune pathogenesis than *P. gingivalis*.

In addition to the absence of specific *P. gingivalis* virulence factors such as hemagglutinin and gingipains, our genomic analysis of *P. catoniae* strains revealed a notable lack of amino acid degradation pathways for glutamate, glutamine, histidine, serine, methionine, tryptophan, and melibiose. Unlike *P. gingivalis*, which utilizes these pathways as carbon sources to support its pathogenic lifestyle, the absence of such metabolic capabilities in *P. catoniae* may suggest a non-pathogenic nature. This potential for commensalism is further supported by the lack of other virulence genes commonly associated with *P. gingivalis*, aligning with findings that characterize non-pigmented *Porphyromonas* species, notably *P. catoniae*, as less likely to contribute to disease states (16, 45). Future studies should incorporate *in vivo* and *in vitro* testing to comprehensively assess these bacteria’s biological roles and impacts.

Moreover, our study identified genes associated with acid resistance in *P. catoniae* strains. Experimentally, these strains demonstrated the ability to grow at a low pH of 5.5 on ABA-SB agar, indicating a robust capacity for acid tolerance. This finding aligns with previous descriptions of *P. catoniae* as a lactic acid producer. This attribute contributes to its survival in acidic environments, much like other lactic acid bacteria, including species of *Lactobacillus* (27). This acid resistance capability not only underscores the adaptive strategies of *P. catoniae* to various environmental stresses but also suggests its potential role in maintaining microbial balance within its niche. The ability to produce and withstand acidic conditions may facilitate the colonization and persistence of *P. catoniae* in the human microbiome, particularly in acid-prone sites, which is a characteristic typical of commensal bacteria.

Our investigation into the antibiotic resistance profiles of pulmonary *Porphyromonas* strains highlights key findings. Consistent with prior studies (27, 46), *Porphyromonas* generally exhibits high susceptibility to antibiotics targeting anaerobes, including β-lactams. However, notable exceptions were observed: *P. uenonis* demonstrated resistance to clindamycin, a relatively new observation given that this species was only recently described (47), and *P. catoniae* displayed resistance to vancomycin, as previously reported (27). In the 2 strains of *P. catoniae*, the search for glycopeptide resistance genes in their genome was negative. Gram-negative bacilli are therefore intrinsically resistant to vancomycin, due to the inability of this glycopeptide to cross the outer membrane. While the present dataset includes only six pulmonary strains, expanding this dataset could provide deeper insights into glycopeptide resistance mechanisms within the POTG group. The isolation of these strains from lung samples remains challenging due to their sensitivity to most antibiotics, oxygen and their slow growth rate.

In conclusion, our study did not detect virulence genes typically associated with *P. gingivalis* in non-pigmented *Porphyromonas* strains, except for a putative hemolysin. These findings suggest that non-pigmented POTG, particularly *P. catoniae*, and *P. pasteri*, are endogenous inhabitants of the lung, contributing to homeostasis rather than disease. This distinction between POTG and the pathogenic *P. gingivalis* is crucial for understanding the lung niche’s microbial dynamics and guiding clinical approaches to lung health and disease.

## 1.5 Abbreviations

(ABB): Anaerobe Basal Broth
(ABA-SB): Anaerobe Basal Agar supplemented with 5% sheep blood
(ARGs): Antibiotic resistance genes
(CA-SFM): Committee for Antibiogram of the French Society of Microbiology
(CF): Cystic fibrosis
(EUCAST): European Committee on Antimicrobial Susceptibility Testing
(HMM): Hidden Markov Model
(MCL): Markov Cluster Algorithm
(MALDI-TOF MS): Matrix-assisted laser desorption/ionization time-of-flight mass spectrometry
(MaGe): MicroScope Microbial Genome Annotation and Analysis Platform
(PY): Peptone Yeast
(POTG): *Porphyromonas* other than *gingivalis*
(RA): Rheumatoid arthritis
(WGS): Whole genome sequencing

## 9. Author statements

### 9.3 Ethical approval

The study was approved by the French Ethical Research Committee in March 2018 (2018-A00624-51). All experiments were performed following relevant guidelines and regulations. Informed consent was obtained from all participants and/or their legal guardians.

## 9.4 Acknowledgements

We gratefully acknowledge the support from LABGeM (CEA/Genoscope & CNRS UMR8030), France Génomique, and the French Bioinformatics Institute. These institutions have provided significant assistance as part of the Investissement d’Avenir program, managed by the Agence Nationale pour la Recherche under contracts ANR-10-INBS-09 and ANR-11-INBS-0013. Their contributions were instrumental in facilitating our use of the MicroScope annotation platform, which was essential for the genomic analyses conducted in this study.

## 11. Supplementary information

**Supplementary Table 1. Screened genes of *Porphyromonas* previously identified by Mendez *et al*. and O’Flynn *et al*. (31, 42)**.

**Supplementary Figure 1.**
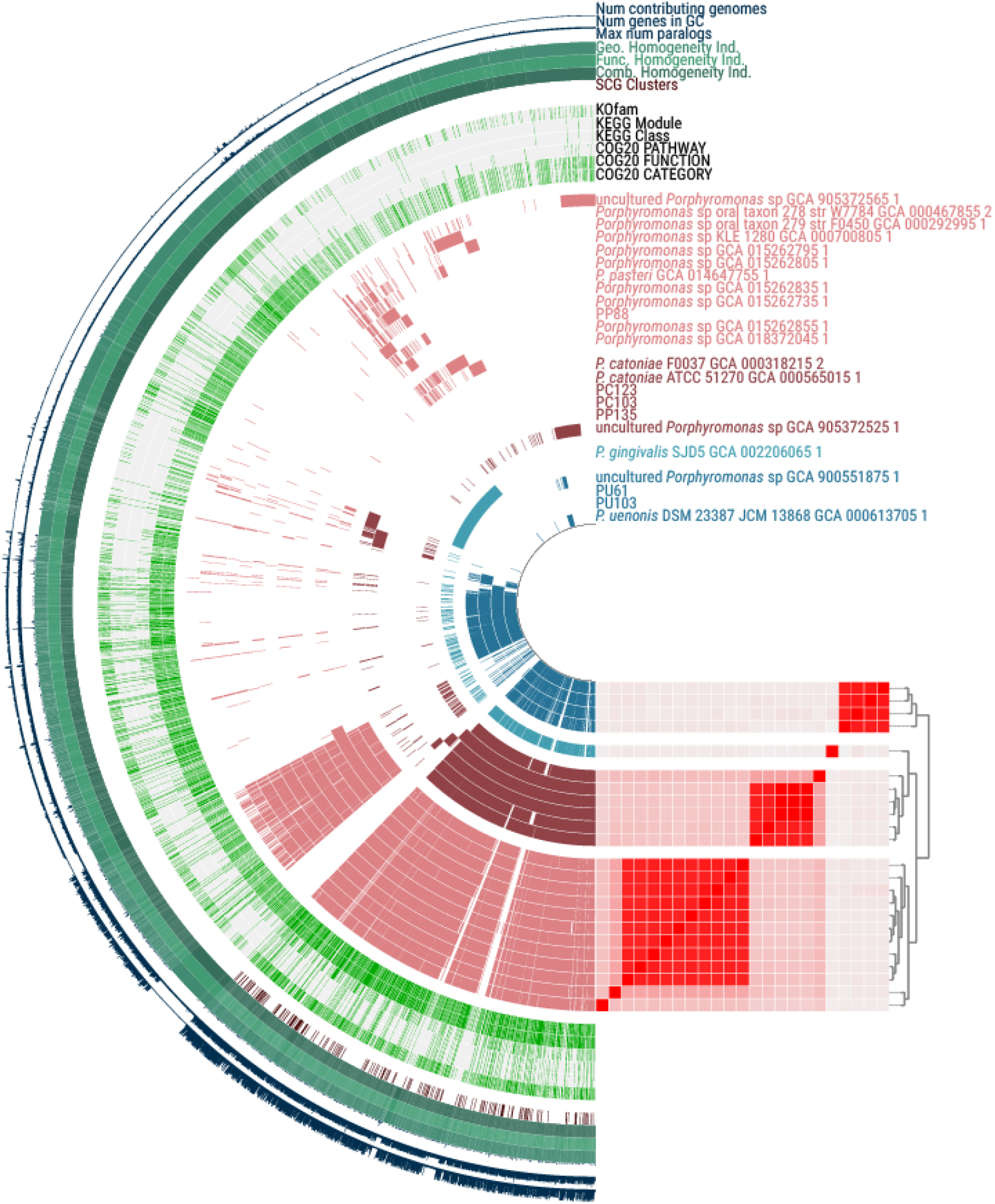
Pan-genome analysis of pulmonary strains of *Porphyromonas*. PC103, PC123, PC135, PP88, PU61, and PU103 were isolated from the lungs of people with cystic fibrosis. Other genomes were downloaded from NCBI.

